# Honeybee flight dynamics and pair separation in windy conditions near the hive entrance

**DOI:** 10.1101/2023.04.14.536844

**Authors:** Bardia Hejazi, Hugo Antigny, Sophia Huellstrunk, Eberhard Bodenschatz

**Affiliations:** Laboratory for Fluid Physics, Pattern Formation and Biocomplexity, Max Planck Institute for Dynamics and Self-Organization (MPI-DS), 37077 Göttingen, Germany; Max Planck University of Twente Center for Complex Fluid Dynamics, MPI-DS, 37077 Göttingen, Germany; Universite Grenoble Alpes, Grenoble INP - Phelma, Grenoble 38000, France; Princeton University, NJ 08544, United States; Institute for Dynamics of Complex Systems, University of Göttingen, 37077 Göttingen, Germany; Laboratory of Atomic and Solid State Physics and Sibley School of Mechanical and Aerospace Engineering, Cornell University, 14853 Ithaca, NY, USA

**Keywords:** Honeybee, Insect, Flight control, Collective behavior

## Abstract

Animals and living organisms are continuously adapting to changes in their environment. How do animals, especially those that are critical to their ecosystem, respond to rapidly changing conditions in their environment? Here, we report on the three-dimensional trajectories of flying honeybees under calm and windy conditions in front of the hive entrance. We also investigate the pitch and yaw in our experiments. We find that the mean velocities, accelerations and angular velocities of honeybees increase with increasing wind speeds. We observed that pair separation between honeybees is highly controlled and independent of wind speeds. Our results on the coordination used by honeybees may have potential applications for coordinated flight of unmanned aerial vehicles.

## 1. Introduction

Animal behavior is influenced by the surrounding environment and changes to the environment. The change in air currents and surface winds impacts the behavior of flying animals, and with the potential increase in wind speeds [1] due to climate change, flying animals may need to adapt their behavior to new conditions. Flying insects such as honeybees are able to adapt to changes in surrounding air flows [2] and maintain flight control even under strongly turbulent conditions [3]. In this work we further explore how honeybees behave and fly in different conditions to better understand their flight dynamics and responses especially when flying in close proximity to one another. The ability for insects to maintain stability and control while in flight [4] and understanding the strategies they use, can aid in improving the systems required for the navigation of small-scale flying robotic collectives [5, 6].

Insects are particularly well suited for research because of their small size, abundance, and the relative ease in which experiments can be carried out on them. Which is why fruit flies have been a long favored example of flying insects for studies [7, 8, 9, 10]. Insects also provide us with the exceptional opportunity to study collective behavior, i.e., their group and social dynamics, as demonstrated by studies of swarming midges [11, 12]. Another insect of particular interest is the honeybee (and other species of bees), due to their important ecological role as natural pollinators [13, 14] on the one hand, but also due to their unique social structure. Experiments studying the social behavior and dynamics of honeybees mostly focus on activity at the hive entrance [15, 16, 17, 18, 19]. Studies that investigate swarms look at stationary collectives that are not in flight, such as work investigating the mechanical stability of honeybee swarms [20, 21].

Experiments studying honeybee flight are usually carried out in the laboratory and investigate the flight of individuals, focusing on wing aerodynamics [22, 23] and dynamical responses to different air flows [2, 24]. Although not as numerous as laboratory studies, field experiments have also investigated honeybee flight stability in windy conditions and observe that honeybees increase roll stability by extending their hind legs [25]. More recently, experiments have studied honeybee flight dynamics outside and looking at the trajectories of honeybees near the hive entrance [26], while also considering the influence of turbulence on honeybee flight [3]. However, there is still a lack of research on the collective flight of honeybees and how they avoid collisions with one another mid-flight. Studies suggest that collision avoidance is achieved through visual sensing [27]. Honeybees may also use other senses for navigation and collision avoidance, for instance, it has been shown that honeybees are capable of sensing electric fields [28, 29] and use this to their advantage when foraging [30] while also contributing to atmospheric electric fields when swarming [31]. In this study we examine the flight dynamics and pair statistics of honeybees near the hive entrance, outside in the open where they are free to fly unhindered. We track honeybees using high-speed, high-resolution cameras and investigate their flight and behavior in natural calm weather conditions and compare to conditions where we generate winds with different mean flow velocities.

## 2. Methods

The experiments were performed at the Max Planck institute for Dynamics and Self-Organization (MPI-DS) in Göttingen, Germany. Experiments were performed on July 27, 2022, from 10:00 to 16:00. The ambient temperature during the experiments was between 20-23 degrees Celsius with no precipitation. Experiments were performed at the entrance of a honeybee hive from the *Apis mellifera carnica* species, as shown in Fig. 1. Performing experiments in front of the entrance allows us to study landing and take-off dynamics and to additionally examine the interactions between pairs of honeybees mid-flight.

**Figure 1.**
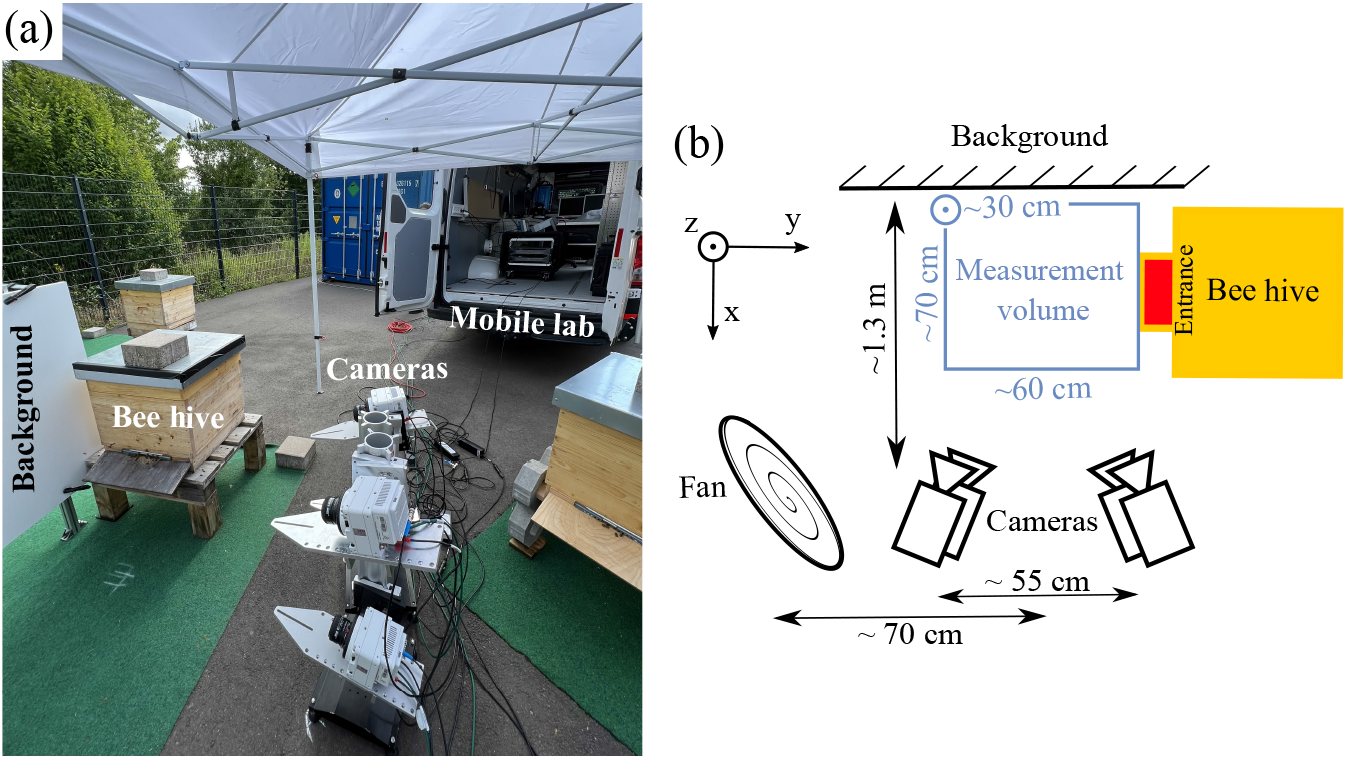
(a) Experimental setup showing the honeybee hive, cameras and background with the overhead shade from the pavilion to protect the cameras from the elements (fan not shown). Also shown is the mobile laboratory where data servers are located and experimentalists monitor recordings. (b) Schematic of the experimental setup showing the location and distances between cameras, background, and fan. The measurement volume in our experiments is approximately 60 (L) x 70 (W) x 30 (H) centimeters where the length is along *ŷ*, the width along 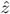, and the height along 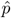.

We created wind in front of the hive using a Trotec model TTV 4500 industrial grade fan with a diameter of 42 cm that had three speed settings. The fan generated a uniform and steady supply of air flow where the vector perpendicular to the plane of the fan was approximately at a 45° angle with respect to − 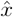 and *ŷ* with a slight upwards tilt in the 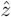 direction. Since the landing platform at the hive entrance was elevated from the ground, the slight upward tilt in 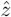 allowed for the generated wind to cover the center of our measurement volume. We measured the ambient air flow with no generated wind and the wind speed produced by the fan using a testo 405i thermal anemometer probe with a resolution of 0.01 m/s. We measured air flow velocities by placing the probe in the center of the measurement volume before each experiment and removing it before recording videos. We performed experiments under three different wind conditions; no wind (with natural air flows), moderate wind (medium wind speed on the fan), and strong wind (maximum wind speed on the fan). The details of the experimental conditions and wind velocities are given in Table 1. The natural air flows were approximately 9% of the moderate and 5% of the strong wind settings of the fan. Thus, the natural air flow was much weaker than the flow produced by the fan.

**Table 1.** Air flow speed measurements along with the number of video recordings for each experiment setting. The mean flow velocity in the x, y, and z direction is reported, with the total mean flow velocity being *U*. The anemometer accuracy as stated by the manufacturer is ±0.1 m/s for speeds from 0 to 2 m/s and ±0.3 m/s for speeds from 2 to 15 m/s.

| Exp. | $U_x$ (m/s) | $U_y$ (m/s) | $U_z$ (m/s) | $U$ (m/s) | No. recordings |
| --- | --- | --- | --- | --- | --- |
| No wind | -0.2 | 0.1 | 0.05 | 0.23 | 25 |
| Moderate wind | -1.0 | 2.2 | 0.5 | 2.46 | 25 |
| Strong wind | -1.9 | 4.0 | 0.6 | 4.46 | 10 |

We recorded images using 4 Phantom VEO4K 990L cameras equipped with 50 mm lenses capturing 4096 × 2034 pixel images at 200 frames per second with approximately 30 seconds of continuous image capture before the camera RAM was full and data transfer was initiated. We recorded a total of 25 videos for the no wind and moderate wind conditions and recorded 10 videos for the strong wind conditions. We recorded less videos for strong wind conditions to limit the time we exposed the honeybees to intense winds and to minimize disturbance to the hive. The cameras had different viewing angles which created an effective measurement volume of 60 (L) x 70 (W) x 30 (H) centimeters. Cameras are triggered and synced using an external signal generator. We used a small pavilion that covered the cameras to protect them from overheating due to direct exposure to the sun and potential harm from any precipitation that may occur during experimentation. We use two data servers for rapid data transfer of video recordings from the camera RAM. The data servers were mounted on a shockproof rack located in a mobile laboratory. The mobile laboratory is a custom van meant for experiments in the field. The experimentalists were seated in the van while experiments were underway and monitored the image capture and download.

Imaging and tracking of honeybees is based on previous methods and in-house codes used to track honeybees using consumer-grade GoPro cameras [3] and similar high-speed Phantom cameras used to track particles in turbulence experiments [32, 33].

The high-resolution cameras used in these experiments allowed us to clearly image honeybees and all their features. From our images we are able to determine honeybee body orientations using a nonlinear fitting algorithm. The routine optimizes for the honeybee position and orientation by minimizing the difference between a model that is projected onto the camera image plane and the actual honeybee images [32]. An example of the orientation finding process is shown in Fig. 2. Fig. 2 (a-d) shows the raw images obtained with our four cameras at the same instance in time. The honeybee identified with the green square frame is seen on all four cameras at this frame. Fig. 2 (e-h) shows the zoomed in image of the honeybee identified in Fig. 2 (a-d). We calculated the honeybee orientation using a 3D capsule-like shape. The raw images were background subtracted and inverted resulting in a clear image intensity gradient from honeybee to background. The modeled shape was also given an intensity gradient similar to the actual image of the honeybee for improved optimization. The 3D model was then projected onto the four two-dimensional (2D) image planes and optimized so that the differences in pixel intensity between the actual honeybee image and projected model image was minimized. With this method we were able to find the honeybee orientation as shown in Fig. 2 (i-l), where the beginning and end of the projected model image are shown on the actual honeybee image. The honeybee center and reprojected center is also shown to demonstrate the tracking precision. The reprojected center was obtained by reprojecting the 3D stereomatched point, obtained using the honeybee center initially found from the four camera images, back to the 2D camera image plane. Fig. 2 (m-p) show the optimized projection of the model capsule-like shape on the 2D camera image planes at the location of the honeybee identified in the previous figure panes.

**Figure 2.**
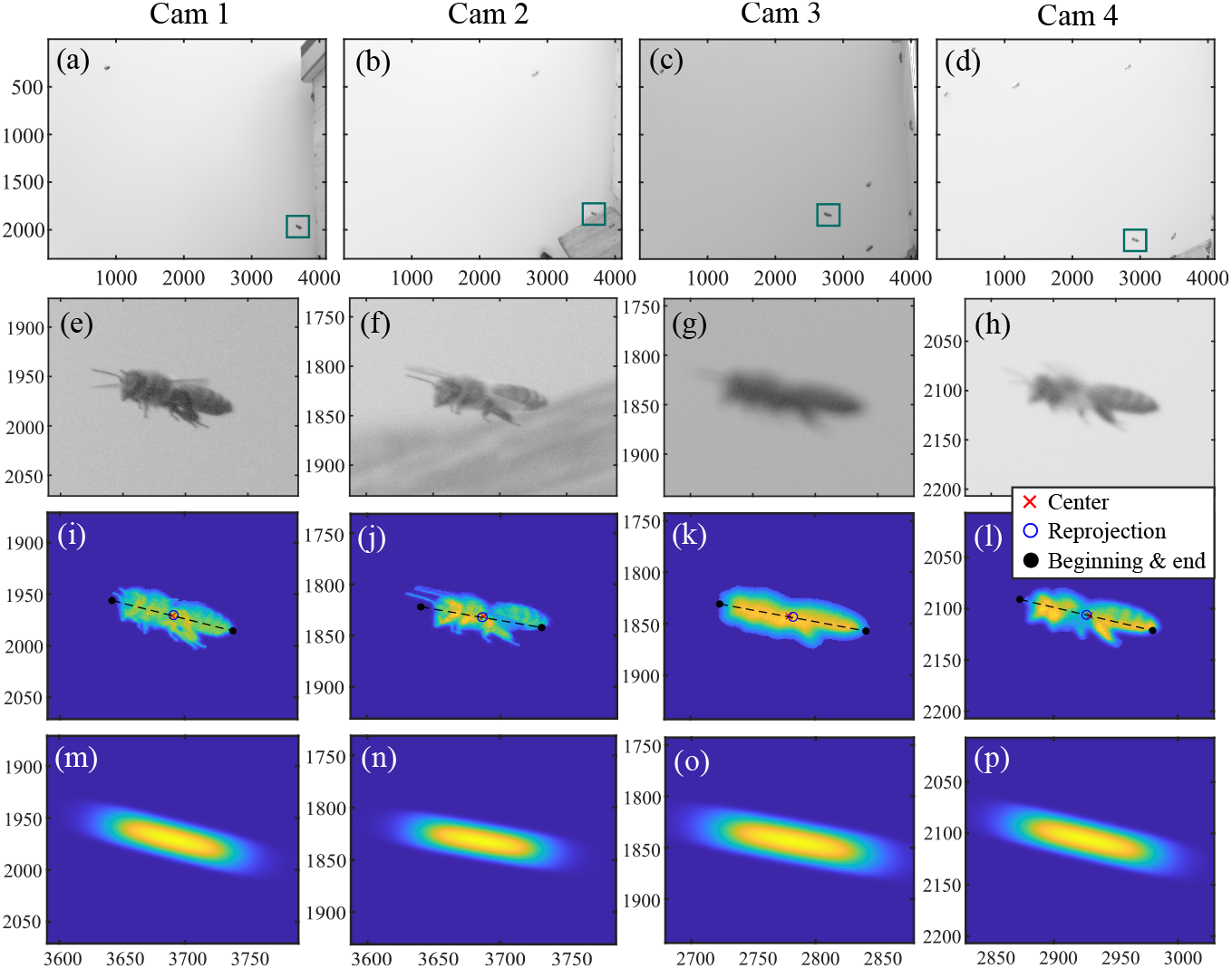
Example of honeybee orientation finding procedure. (a-d) raw images from the cameras at the same frame number where an identified honeybee is seen on all four cameras and is tracked. (e-h) Zoomed in image of the identified honeybee in (a-d). (i-l) Background subtracted and processed image of the honeybee where the center, beginning, and end of the honeybee are identified by fitting a capsule-like shape to the honeybee. (m-p) The capsule model that is fitted to the honeybee at the same position as the honeybee in (i-l).

## 3. Results

We calculated honeybee flight velocities and accelerations using polynomial fits to the three-dimensional (3D) trajectories with fit-length of *h* = 7 where each track of length *l* is split into *l* − 2*h* sections for fitting. As a result, we only analyzed tracks longer than 10 frames or 50 ms. Furthermore, we only used data in the analysis where the stereomatching error was no more than 5 pixels. The resulting mean tracking error across all cameras was approximately one pixel where an individual bee is approximately 4000 pixels in size. Table 2 shows the total number of tracks, mean track length, and maximum track length of each wind condition experiment. We can see that as the wind intensity increases the mean and maximum track lengths decreases, thus honeybees seem to not linger in windy conditions when compared to mild, no wind conditions.

**Table 2.** The total number of tracks, mean track length, and maximum track length for each experiment setting.

| Exp. | No. tracks | Mean track length | Max. track length |
| --- | --- | --- | --- |
| No wind | 3894 | 59 | 504 |
| Moderate wind | 3948 | 43 | 459 |
| Strong wind | 797 | 37 | 272 |

Figure 3 shows the probability distribution function (PDF) of the honeybee velocity and acceleration components in different wind conditions. From Fig. 3 (a,c,e) we observe an unsymmetrical pattern in the velocity component PDF’s. Here when there is no wind generated by the fan (blue filled circles), the honeybee has larger velocities when leaving the hive as shown by the skewed PDF of *v*_*y*_ towards larger negative values. While, the other two components of *v*_*x*_ and *v*_*z*_ remain symmetric. However, as we introduced wind, the velocity component PDF’s changed in that the honeybee velocity is skewed in the same direction as the mean wind velocity for all components of *v*. The acceleration components do not strong signs of asymmetry but as the wind intensity increases, the mean acceleration magnitude also increases, this is also the case for the mean honeybee velocity magnitude. The mean honeybee velocity and acceleration magnitudes in different wind conditions are given in Table 3.

**Table 3.**
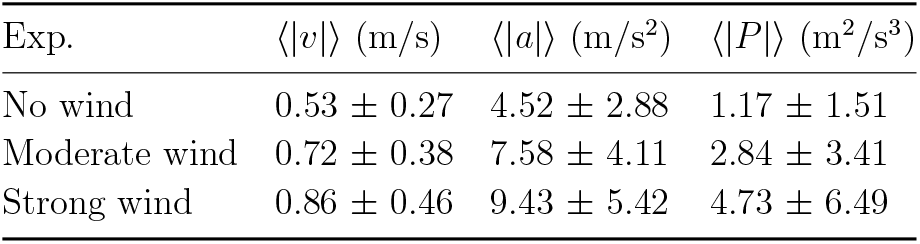
The mean velocity, acceleration, and power magnitudes for each wind condition experiment ± the standard deviation.

**Figure 3.**
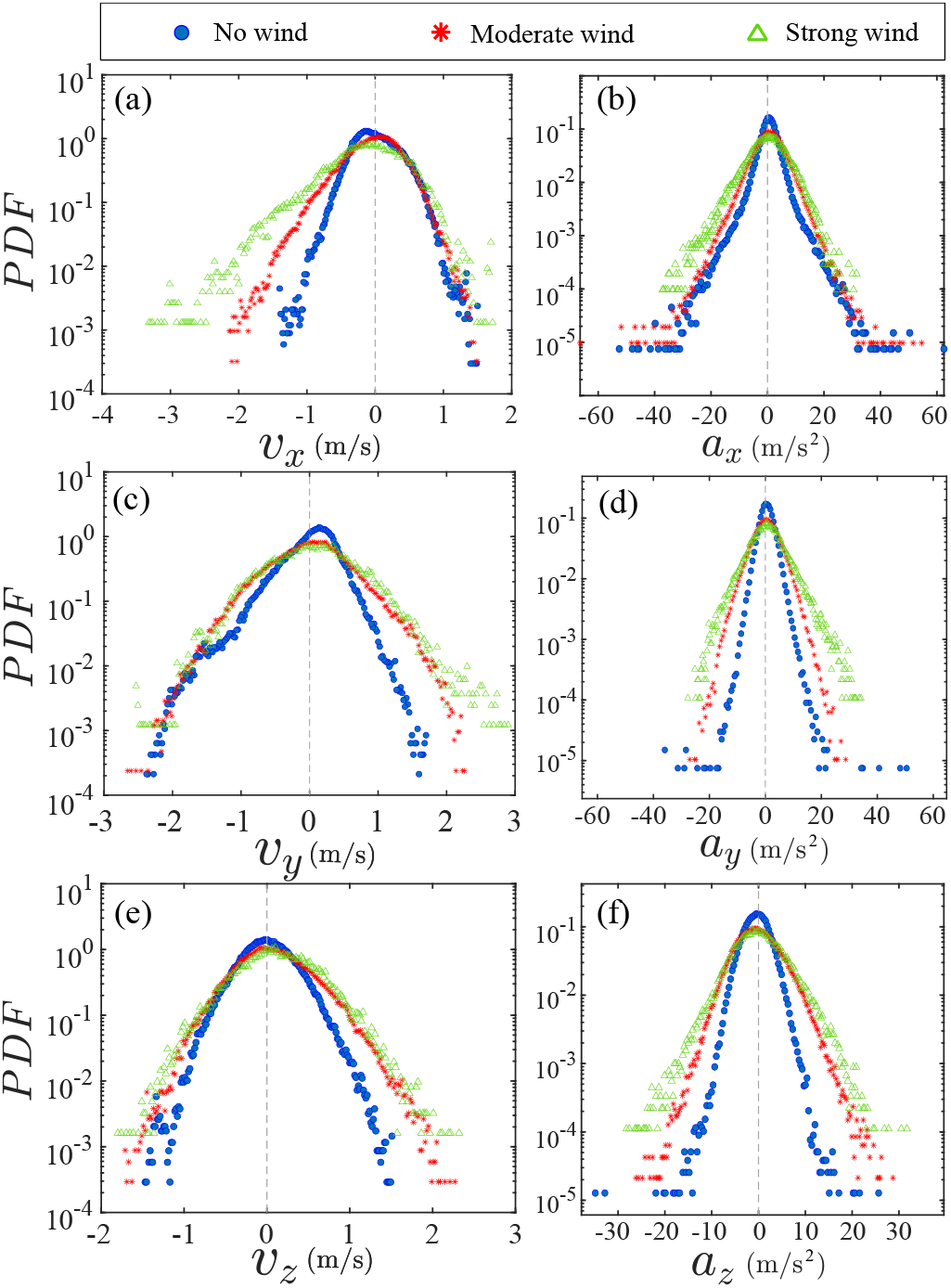
The probability distribution function (PDF) for honeybee velocity and acceleration components in different wind conditions. (a,b) *x* components, (c,d) *y* components, and (e,f) *z* components. The mean velocity and accelerations increase with wind strength. The mean wind direction is in the +*y*, −*x*, and +*z* direction.

Figure 4 shows the PDF of the honeybee flight power per insect mass, defined as *P* = *a* · *v*. We assume that the mass difference between honeybees is negligible as compared to the variance in their velocity and acceleration. We calculate the flight power using definitions from Lagrangian turbulence used to determine the change in kinematic energy of tracer particles advected in turbulent flow [34]. Similarly to the mean velocity and acceleration magnitudes, the mean flight power magnitude also increases with increased wind velocity. In previous work we had observed that the mean flight performance of honeybees (⟨|*v*|⟩, ⟨|*a*|⟩, and ⟨|*P* |⟩) did not change as we changed the wind turbulence intensity [3]. In our previous experiments the mean wind velocity did not change significantly and was approximately uniform for all turbulence intensities investigated. However, now by significantly changing the mean wind velocity we clearly see a dependence of the flight performance on wind intensity.

**Figure 4.**
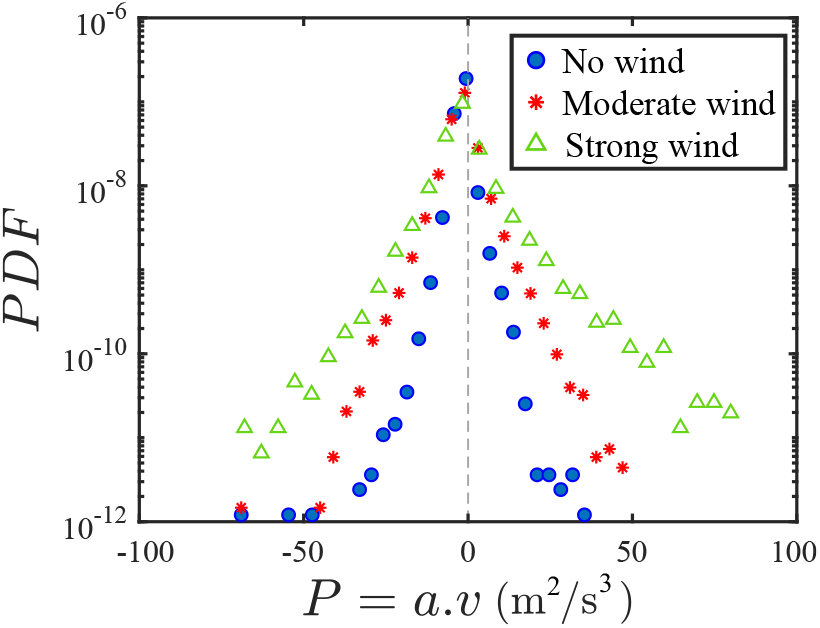
The PDF of honeybee flight power, *P* = *a* · *v*, for different wind conditions. The mean flight power is increasing with mean wind velocity.

Performing experiments in front of the hive entrance allowed us to also study the differences between landing and takeoff dynamics. For example, we showed that in no wind conditions (Fig. 3 (a)), honeybees have larger velocities when leaving the hive.

Figure 5 shows the cosine of the angle between the honeybee body vector, 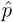, and the horizontal axis *ŷ*. The vector that defines the honeybee body orientation is the vector that lies along the length of the honeybee body from the tail to head or end to beginning of the honeybee as shown in Fig. 2 (i-l). In Fig. 5 (a), 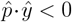 and the honeybee is facing away from the hive and is generally flying away from the hive. Similarly in Fig. 5 (b), 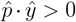 and the honeybee is facing towards the hive and is generally flying towards the hive. The reason we use the term generally is that occasionally when the hive entrance was crowded the honeybees hovered or flew backwards away from the hive while facing towards the hive. The honeybees seemingly have a preferential body orientation while they fly away and towards the hive in no wind conditions, as can be seen by the maxima in the PDF of the cosines. In no wind conditions, the honeybees have larger angles away from the horizontal as they approach the hive in anticipation of landing as compared to when leaving the hive with larger velocities (Fig. 3 (c)). However, this preferential body angle while flying disappears when we introduce wind and the honeybees prefer to fly in a way that their body is mostly horizontal and aligned with *ŷ*. Further investigation is needed to determine why a more horizontal flight is preferred in windy conditions and if it causes increased flight stability in windy and unstable conditions.

**Figure 5.**
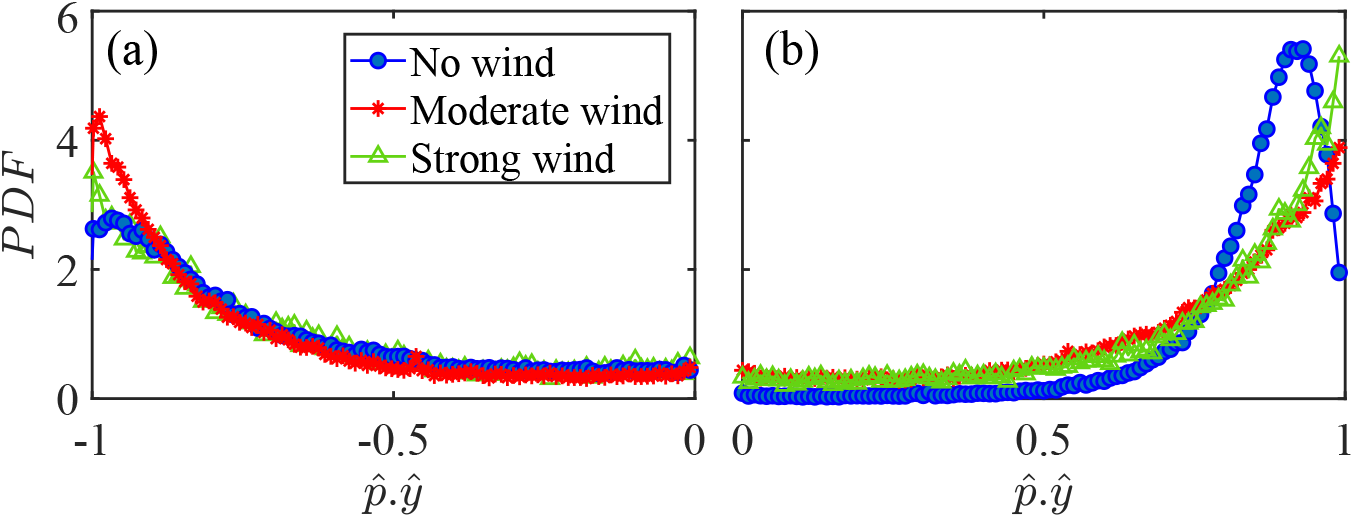
Cosine of the angle between the honeybee body vector, 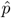, and the global *ŷ* vector for different wind conditions. (a) Honeybee is facing away from the hive and is generally moving away from the hive, (b) honeybee is facing towards the hive and is generally moving towards the hive.

We calculated the honeybee angular velocity from the honeybee body orientation. Fig. 6 shows the PDF of the honeybee angular velocity where *ω*_*n*_ = (1*/*3)(*P* (*ω*_*x*_*/ω*_*x*_′) + *P* (*ω*_*y*_*/ω*_*y*_′) +*P* (*ω*_*z*_*/ω*_*z*_′)), and *ω*_*i*_′ is the standard deviation of *ω*_*i*_. The angular velocity does not represent the complete angular dynamics of the honeybee since with our orientation finding method we can only obtain two of the three flight angles. The angular velocity shown in Fig. 6 represents the changes in the pitch and yaw of the honeybee and does not include roll dynamics. Even so, we can still observe interesting dynamics where just as before for the other flight parameters, the mean angular velocity increases with wind mean flow velocity. Table 4 shows the values for the mean angular velocity magnitudes for each wind condition experiment.

**Table 4.**
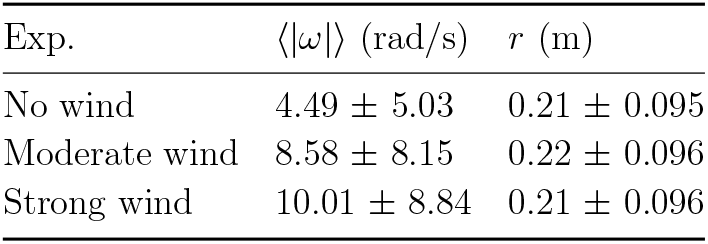
The mean angular velocity magnitudes and pair separations for each wind condition experiment ± the standard deviation.

**Figure 6.**
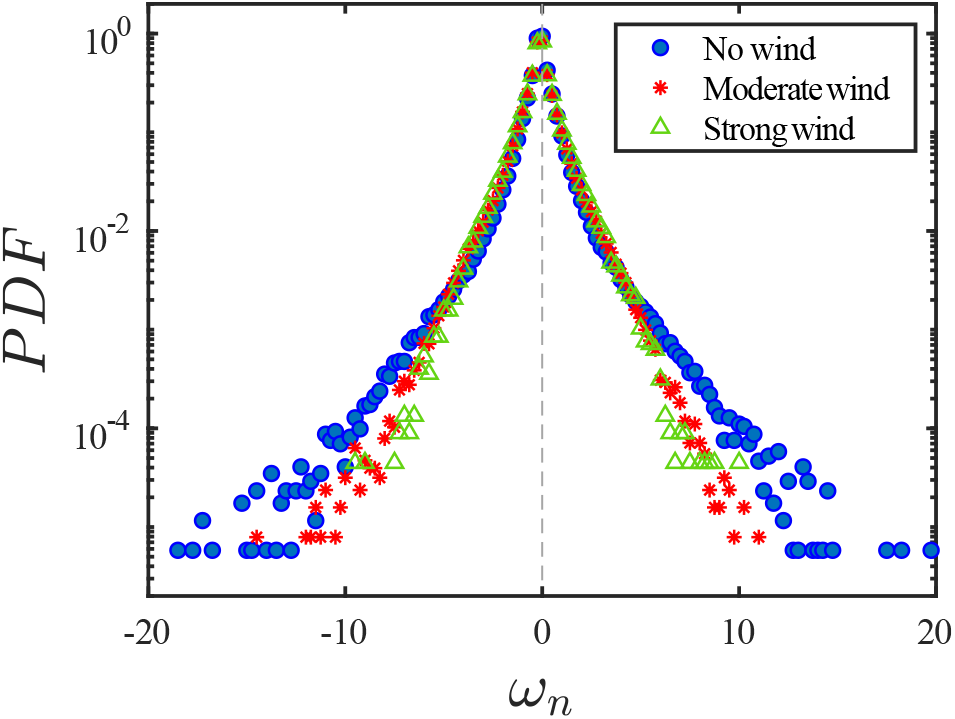
The PDF of honeybee angular velocity in different wind conditions. Here the angular velocity is calculated based on the pitch and yaw angles. The mean of angular velocities increases with mean wind velocity.

An advantage of performing experiments in front of the hive entrance was that we were able to image multiple honeybees simultaneously, from at least two to upwards of ten or more honeybees in the same frame. This allowed us to obtain pair statistics such as the PDF of the distance between every two honeybees seen at the same instance in time, Fig. 7 shows this PDF and Table 4 shows the mean pair separations for all wind conditions. The honeybees appear to be maintaining a constant distance from one another independent of the wind intensity. While the honeybees seemingly have less control in their flight in extreme conditions since all the other flight parameters (*v, a, P*, and *ω*) were affected by the change in wind intensity, they still maintain a chosen distance from one another.

**Figure 7.**
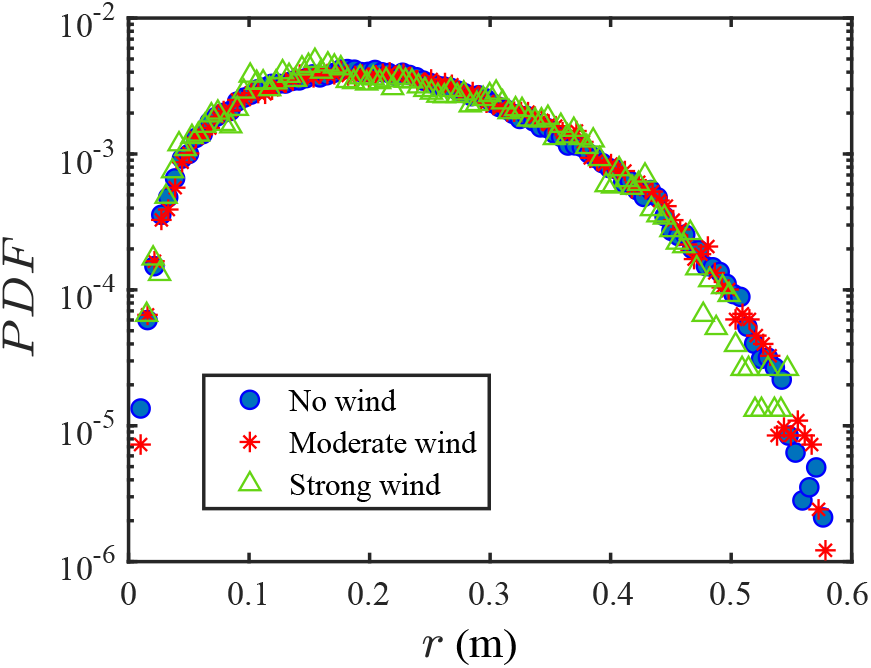
The PDF of the distance between every pair of honeybees seen at the same instance in time for different wind conditions. Honeybees display the same behavior in all wind conditions.

## 4. Conclusions

The mean honeybee velocity, acceleration, flight power, and angular velocity were all influenced by the mean wind velocity where the mean magnitude of all the quantities measured increased with the mean wind velocity. In previous work we had observed at constant mean wind magnitudes that the measured quantities were not dependent on the turbulence intensities [3]. Thus we conclude that it is the mean wind and not the turbulence intensity to which the honey bees respond by changing these properties. In calm no wind conditions the honeybees have a preferential orientation in which they approach and leave the hive. The approach angle is larger than the leaving angle. With wind the honey bees align their bodies with the horizontal axis. Such behavior is perhaps a way to increase flight stability in windy conditions. The limitation to our orientation finding method is that we can only identify two of the three flight angles, namely we can only measure changes in the pitch and yaw. In future work we will also measure roll dynamics with the use of more advanced computer vision techniques [35]. By performing experiments in front of the hive entrance we explored the interaction between individual honeybees. We found that the honeybees maintain a constant distance from one another during flight for all wind conditions. This distance was kept approximately constant even under extreme wind conditions while the honeybees had to adjust their flight parameters. Future work would need to explore this phenomena in more detail and explore under what conditions the pair separation between honeybees can be influenced, such as introducing electric fields and whether such an external field disrupts navigation and collision avoidance. The results presented here on how honeybees fly in extreme winds and how their flight is affected, from their body orientation to their pair separation, can be of great interest to the advancement of flight stability, control, and the systems needed for collision avoidance in collective robotic motion.

## Acknowledgments

This work would not have been possible without the support of the technical staff at the MPI-DS. The authors would like to thank the biology staff for taking care of the honeybee hives. We would like to thank the experimental technicians for help with setting up the mobile laboratory and pavilion. Additionally, many thanks to the IT and HPC team for the setup of the data servers for file transfer and storage. We would like to express our sincere gratitude to Alain Pumir for insightful discussions.

## Data and code availability

Available upon request.

## Author contributions

BH planning and development of research project. BH, HA, and SH performing experiments and initial orientation analysis. BH analysis of data and writing original draft. BH, HA, SH, and EB editing and preparation of final draft.

## Ethics statement

According to the representative for animal experiments in basic research at the Max Planck Society, no ethical or legal statements are required for the experiments presented in this work.

